# A novel inovirus reprograms metabolism and motility of marine Alteromonas

**DOI:** 10.1101/2022.06.23.497286

**Authors:** Kuntong Jia, Yongyi Peng, Xueji Chen, Huahua Jian, Min Jin, Zhiwei Yi, Ming Su, Xiyang Dong, Meisheng Yi

**Author notes:** Correspondence: Meisheng Yi: School of Marine Sciences, Sun Yat-sen University, Guangzhou 510275, Guangdong, China, Xiyang Dong: Key Laboratory of Marine Genetic Resources, Third Institute of Oceanography, Ministry of Natural Resources, Xiamen 361005, Fujian, China.

## Abstract

Members from the *Inoviridae* family with striking features are widespread, highly diverse and ecologically pervasive across multiple hosts and environments; however, very small amount of inoviruses have been isolated and studied. Here, a filamentous phage infecting *Alteromonas abrolhosensis*, designated ϕAFP1, was isolated from the South China Sea and represented as a novel genus of *Inoviridae*. ϕAFP1 consisted of a single-stranded DNA genome (5986 bp), encoding eight putative ORFs. Comparative analyses revealed ϕAFP1 could be regarded as genetic mosaics, which especially came from *Ralstonia* and *Stenotrophomonas* phages. The temporal transcriptome analysis of *A. abrolhosensis* to ϕAFP1 infection reveals that 7.78% of the host genes were differentially expressed. The genes involved in translation processes, ribosome pathways and degradation of multiple amino acid pathways at plateau period were upregulated, while host material catabolic and bacterial motility-related genes were downregulated, indicating that ϕAFP1 might hijack the energy of the host for the synthesis of phage proteins. ϕAFP1 exerted the step-by-step control on host genes through the appropriate level of the utilizing host resources, affirming a new non-standard regulatory strategy of viral temperately control over the host transcriptional profile. Our study provides novel information for a better understanding of filamentous phage characteristics and phage-host interactions.

## Introduction

Bacteriophages, the viruses of bacteria, are the most abundant biological entities in the global ecosystem. The complex, dynamic phage-bacteria interactions make phages have dramatic influences on the diversity and physiology of their hosts, the marine nutrient cycle and horizontal gene transfer (1, 2). Most of the known phages belong to the order of *Caudovirales* which are tailed phages with an isometric capsid and divided into three phylogenetically-related families: *Myoviridae*, *Siphoviridae*, and *Podoviridae*. Only a few phages accounting for less than 4% of all known phages are shaped as polyhedral, filamentous, or pleomorphic (3). Among them, filamentous bacteriophages belonging to *Inoviridae*, *Lipothrixviridae*, and *Rudiviridae* families, are relatively rare (4, 5).

All phages from the *Inoviridae* family (inoviruses) share a similar filamentous virion shape which is typically about 6-8 nm in width and 800-2000 nm in length containing a circular single-stranded DNA (ssDNA) genome of nearly 4-12 kb (6–8). The genome of filamentous phage encodes around 10 genes which compose of four functional modules involved in genome replication, virion structure, assembly/secretion, and regulation (9). Almost all of the described filamentous phages were isolated from Gram-negative bacteria (10), while there were only two filamentous phages that were found in the Gram-positive bacteria, *Propionibacterium freudenreichii* and *Clostridium acetobutylicum* (11). Unlike the typical bacteriolytic tailed phages, filamentous phages can produce new mature virions which are even packaged on the cell surface and then secreted to the extracellular space without killing the host bacteria (12). In addition, like many other phages, filamentous phages can undertake wide-ranging DNA recombination to catch new genes and perform as important shuttles for gene transfer among host cells, which may promote bacterial evolution (13, 14).

The genus *Alteromonas* contains species of marine bacteria that are described as Gram-negative, strictly aerobic, and globally oceanic distributed heterotrophic bacteria with an unsheathed single polar flagellum (15). Members of this genus are widely found in surface seawater, open deep ocean, and coastal seawater (16, 17). Despite *Alteromonas* strains are virtually ubiquitous and play a crucial role in the biogeochemical cycling of marine ecosystems, the knowledge regarding the phage-host interaction is still poor, especially the impact of temperate phages infecting *Alteromonas*.

Phages take over host resources for reproduction by manipulating hosts’ critical biological pathways involved in transcription, translation, signal transduction and metabolism (18, 19). However, the relationship between filamentous bacteriophages and their hosts is not only antagonistic, but also mutualistic in many aspects. Several researchers have reported some filamentous bacteriophages can affect the virulence factor and pathogenicity of the host by carrying cholera toxin genes or activating hosts’ virulence genes (20, 21). Moreover, filamentous bacteriophages may be beneficial to hosts’ adaptive strategies in the deep-sea environment, which have been shown to significantly regulate the synthesis of transcription and translation elements, as well as the swarming motility of deep-sea bacteria for improved host survival (22, 23). In addition, *Pseudomonas aeruginosa* phage Pf1 has been known to enhance the resistance of biofilm by their hosts to environmental stress (24).

Relative to the abundant existence of *Alteromonas* strains in the marine environment, the sparsely reported phages infecting *Alteromonas* showed a huge contrast. In this work, we isolated and purified a new filamentous phage ϕAFP1 that infects *Alteromonas abrolhosensis* strain from a seawater sample collected from the South China Sea, and characterized its morphology, biological characteristics, genomic features and gene structure-function analysis. We further investigated the first global glimpse of the interactions between *Inoviridae* and *Alteromonas* by a combination of whole-genome sequencing, RNA sequencing (RNA-seq) and functional assay. This study expands our understanding of *Alteromonas* phage diversity in the marine environment, together with the genomic characteristics and phage-bacterium interaction of filamentous phages.

## Materials and methods

### Bacterial strains and culture conditions

The bacterial strains (*Pseudomonas putida*, *Pseudomonas mosselii*, *Pseudomonas sp*, *Vibrio* strain) used for host range test were isolated and kept in our laboratory. *Edwardsiella tarda* strain (ATCC 15947) was bought from American Type Culture Collection (ATCC). The host *A. abrolhosensis* strain JKT1 was sampling near the beach of Tangjia Bay (Zhuhai, Guangdong, China). All the strains were grown at 28°C on 2216E marine agar medium (Hopebio, China).

### Phage isolation, purification and concentration

The seawater sample used for ϕAFP1 isolation was collected from the northern sea area of Xisha Islands (Paracel Islands) in the South China Sea (17°04′37.27″N, 111°28′24.87″E) during the summer of 2019. The phage isolation assay was performed as described previously with some modification (25). The seawater sample (5 ml) was added to an equal volume of 2216E liquid medium and inoculated with 750 μl of *A. abrolhosensis* strain JKT1 overnight culture in an incubator shaker (28℃) (Crystal, USA) for 18 h. Then, the culture was centrifuged and the supernatant was filtered through a 0.22 μm-pore-size sterile PES syringe filter (Sorfa, China) to remove bacteria cells and debris.

Phage was detected by the soft-agar overlay method with some modifications (26). The filtered solution was incubated with *A. abrolhosensis* strain JKT1 at RT for adsorption, mixed with soft agar and then poured on a bottom agar plate immediately. After overnight cultivation at 28℃, the obvious plaques were picked and suspended in SM buffer. Six rounds of plaque isolation experiments were carried out to acquire purified phages, then which were stored in SM buffer at 4℃.

Phage ϕAFP1 was condensed by the standard protocol described by Sambrook and Russel (27). Overnight culture of phage ϕAFP1 and *A. abrolhosensis strain* JKT1 (1 L) was pelleted at 15,000 × g for 10 min. The supernatant was mixed with polyethylene glycol 8000 and bathed in the ice for 2 h. Phage particles were precipitated at 10,000 × g for 10 min at 4°C, after which the pellet was resuspended by SM buffer, following by the addition of Cesium Chloride (0.75 g ml^-1^) (CsCl, Sigma-Aldrich, USA). The mixture was transferred to an ultra-clear centrifuge tube (Beckman Coulter) and centrifuged in an Optima XPN-100 Ultracentrifuge (Beckman Coulter, USA). After which the phage pellet was resuspended with SM buffer and kept at −80°C.

### Morphological observation

One drop of high-titer purified phage ϕAFP1 solution was subjected to carbon-coated copper grids (400 mesh) for adsorption for 2 min before negatively stained with 2% (w v^-1^) phosphotungstate (pH 7.0), then observed with a JEM-1400 electron microscope at a working voltage of 120kV.

### Host range test

The spot test was performed to characterize the host specificity of phage ϕAFP1 according to the previous study with modification (28). Briefly, host bacterial culture (0.1 ml) (*P. putida*, *P. mosselii*, *Pseudomonas sp*, *E. tarda,* and *Vibrio* strain) was mixed thoroughly with 2.5 ml of upper layer molten 2216E agar (0.7%) medium before poured on the surface of a solid 2216E agar (1.5%) plate. After the upper layer was completely solidified, phage suspension (5 μl) was dropped onto the upper layer and incubated for 16 h at 28℃. The sequence-based tools PHIS Detector (http://www.microbiome-bigdata.com/PHISDetector/) was used for detecting phage-host interactions (29).

### Biological characteristics and bacteriolytic activity *in vitro*

Thermal and pH stability, and chloroform sensitivity were tested as described previously (30). For thermal stability, phage suspension (10^8^ pfu ml^-1^) was sustained separately at different temperatures of −20℃, 4℃, 28℃, 37℃, 50℃, 60℃ and 70℃ for 2 h. For pH stability, phage stock (10^8^ pfu ml^-1^) was individually added into SM buffer with different pH values from 2 to 10 and then incubated for 2 h. To assess the influence of chloroform, chloroform was added into an aseptic tube which contained phage stock (10^8^ pfu ml^-1^).

The MOI assay was performed as described previously with some modifications (31). The phage stock solution was diluted in a gradient with SM buffer and then infected *A. abrolhosensis* strain JKT1 (10^8^ cfu ml^-1^) at an MOI of 0.001, 0.01, 0.1, 1, 10 for 20 min at RT, respectively. The MOI value which yielded the maximum phages would be considered as the optimal MOI of ϕAFP1.

### One-Step Growth Curve

A one-step growth curve assay was performed as described previously with some adjustments (32). *A. abrolhosensis* strain JKT1 culture (10 ml, OD_600_=0.4∼0.5) was harvested by centrifugation (2,500 × g) at RT for 10 min. Then the pellet was resuspended in SM buffer and added with phage stock solution (10^9^ pfu ml^-1^) at an MOI of 1. After 20 min’ standing at RT, the mixture was centrifuged (2,500 × g) at RT for 5 min and washed by SM buffer for three times to remove free phage particles. The pelleted mixture was carefully resuspended in 2216E liquid medium, then incubated in a shaker (180 rpm) for 8 h at 28℃. During the cultivation, a phage sample was taken at 3-min intervals for the first 15 min, 15-min intervals for the next 45 min and 1-h intervals for the remaining 7 h, until 8 h. Every sample was immediately centrifuged (8,000 × g) once it was taken, then the supernatant was transferred to a new sterile tube and stored at 4℃ temporarily. Finally, the phage titer was determined by the soft-agar overlay method mentioned above.

### Genomic DNA and protein analysis

The genomic DNA of phage ϕAFP1 was extracted using a Viral DNA Kit and separated on 1% (w v^-1^) agarose gels. For analyzing phage genomic DNA characteristics, phage DNA was treated with DNase I, RNase A, S1 nuclease and general restriction enzymes.

Tris-tricine-SDS-PAGE was carried out to analyze structural proteins of phage ϕAFP1 (33). The purified phage particles (10^12^ pfu ml^-1^) were mixed with 2 × lysis loading buffer (Solarbio, China) and boiled for 10 min. Thereafter, the prepared sample was separated by electrophoresis on a 20% Tricine-SDS polyacrylamide gel (Solarbio, China) which was then stained with Coomassie brilliant blue R-250 (Amresco, USA) to visualize.

### Sequencing and assembly of phage ϕAFP1

Whole-genome sequencing of ϕAFP1 was performed using a Truseq SBS Kit (300 cycles, Illumina) on Illumina NovaSeq 6000 (Illumina Inc.) at LC Bio (Zhejiang, China). The original sequencing data of phage ϕAFP1 were processed and trimmed by fastp v.0.12.4 to obtain high-quality clean reads (34). Potential viral contigs were assembled by MetaViralSPAdes v.3.15.2 pipeline with default parameters (35). CheckV was used to evaluate the completeness and contamination of assembled genomes of phage ϕAFP1 (36).

### Sequencing and assembly of *A. abrolhosensis* strain JKT1

The genomic DNA of *A. abrolhosensis* strain JKT1 was extracted by Bacterial DNA Kit D3350 according to the manufacturer’s instructions. The genomes of *A. abrolhosensis* were sequenced using the PacBio RS II and Illumina NovaSeq platforms. Raw Illumina sequencing reads were trimmed, filtered and quality screened using FastQC v.0.11.8. The assembly of the whole-genome sequence was conducted by Unicycler v.0.4.8, a hybrid assembly pipeline for bacterial genomes (37). Completeness of assembled bacterial genomes was estimated using CheckM (38). The corresponding taxonomy of the bacterial host was assigned by GTDB-TK v202 with reference to GTDB R06-RS202 and classified as *A. abrolhosensis* strain JKT1 (39).

### Genome annotation

GeneMarkS-2 (http://opal.biology.gatech.edu/GeneMark/index.html) was used to predict the encoding genes of the assembled genomic sequence of phage ϕAFP1 (40). Functional annotation of *A. abrolhosensis* strain JKT1 was performed using EggNOG-mapper v.2.1.7 (41) based on eggNOG orthology data (42) and sequence searches were performed using DIAMOND (43).

### Phylogenetic analysis and network analysis

Whole-genome sequences-based phylogenetic analysis was performed and visualized among all 38 related filamentous phages in GenBank (Supplementary Table 4) using ViPTree (44). The sequence similarities of the selected genomes are calculated by ViPTree according to tblastx.

The network analysis of viral genomes implemented in vConTACT v.2.0 with default parameters to identify genus-level groupings of phage ϕAFP1 (45). The clustered viruses were able to be assigned to the related viral genus and family level in Viral RefSeq v201 reference database.

### Comparative genomic analysis and structural modelling

Assembled viral genomes were compared with reference genomes and visualized using EasyFig v2.2.2 with the tBLASTx option and the filtering of small hits/annotations option (E-value cutoff of 1×10^−3^ and min. length of 15 residues) (46). Structural modeling for all was performed using RoseTTAFold (47). Protein structures were rendered by PyMoL v.2.5.2.

### Transcriptomics

Samples (three biological replicates) for transcriptomics were taken in the three time points (6 min, 2 h, and 6 h) for latent, burst and plateau periods of its one-step growth curve. The cultivation of *A. abrolhosensis* strain JKT1 and ϕAFP1 was performed as described above. Then the supernatant was decanted, and precipitation was collected. RNA was extracted and quantitated with a Qubit 2.0 fluorometer. The rRNA depletion with the Epicentre Ribo-Zero rRNA Removal Kit, library construction with the NEBNext® Ultra II™ Directional RNA Library Prep Kit for Illumina, and paired-end sequencing on an Illumina NovaSeq 6000.

RNaseq reads were quality filtered and trimmed by fastp v.0.12.4, then the ribosomal RNAs in clean data were removed by comparing with rRNA sequences in Rfam and Silva database by SortMeRNA v.4.2.0. The trimmed reads for each library were aligned to the transcripts of *A. abrolhosensis* strain JKT1 and ϕAFP1 using Bowtie2 v.2.3.5. Preprocessed reads were mapped to the *A. abrolhosensis* strain JKT1 genome to generate read count quantification TPM (Transcripts Per Million) of each transcript using Salmon v.1.8.0. DESeq2 v.1.30.1 was used to calculate individual transcript expression multiple and pairwise q-values (FalseDiscovery Rate rectification by Benjaminiand Hochberg method) (48). Fold-change was calculated in Excel as expression level in case / expression level in control. Differentially expressed genes were defined as those p adjust value ≤ 0.05 and |log2(fold-change)| ≥ 1. GO annotation and KEGG enrichment analysis were performed using clusterProfiler v.3.18.1 (49).

### Quantitative reverse transcription-PCR

Quantitative reverse transcription-PCR (qRT-PCR) was performed to validate the RNA-seq data. The cDNA synthesis was performed with GoScript™ Reverse Transcription Mix kit according to the manufacturer’s recommendations. Primers for RT-qPCR are listed in Supplementary Table 14. 16S rRNA gene was used as the reference gene for normalization. RT-PCR was performed using Light Cycler 480Ⅱ real-time PCR system with ChamQ Universal SYBR qPCR Master Mix. The following cycling conditions were used: with 45 cycles consisting of 5 min at 95°C, 10 s at 95°C, 35 s at 60°C and 2 min at 40°C.

### Statistics analysis

All statistical analyses were performed using SPSS (version 19). One-way ANOVA was applied for the assessment of the differences between control and treatment groups. The statistically significant and extremely significant differences were represented by *p* < 0.05 (*) and *p* < 0.01 (**), respectively.

### Data availability

The whole genome of phage ϕAFP1 and *A. abrolhosensis* strain JKT1 was deposited at GenBank under accession number MK473382. The raw sequence of RNA-seq is available in the SRA database under accession number SRR13374388 (BioProject no. PRJNA689384).

## Results

### Morphology and characterization of ϕAFP1

A filamentous phage infecting *A. abrolhosensis* strain JKT1, named ϕAFP1 was isolated from the South China Sea. Phage ϕAFP1 formed turbid, circular and boundary smooth plaques with 0.5-1.0 mm in diameter in a double agar layer at 28°C while no plaque appeared in control (Figure 1A). The morphology of ϕAFP1 was observed by Transmission electron microscopy (TEM) and appeared to be filamentous in shape approximately 10 nm in width and 900 nm in length (Figure 1B).

**Figure 1.**
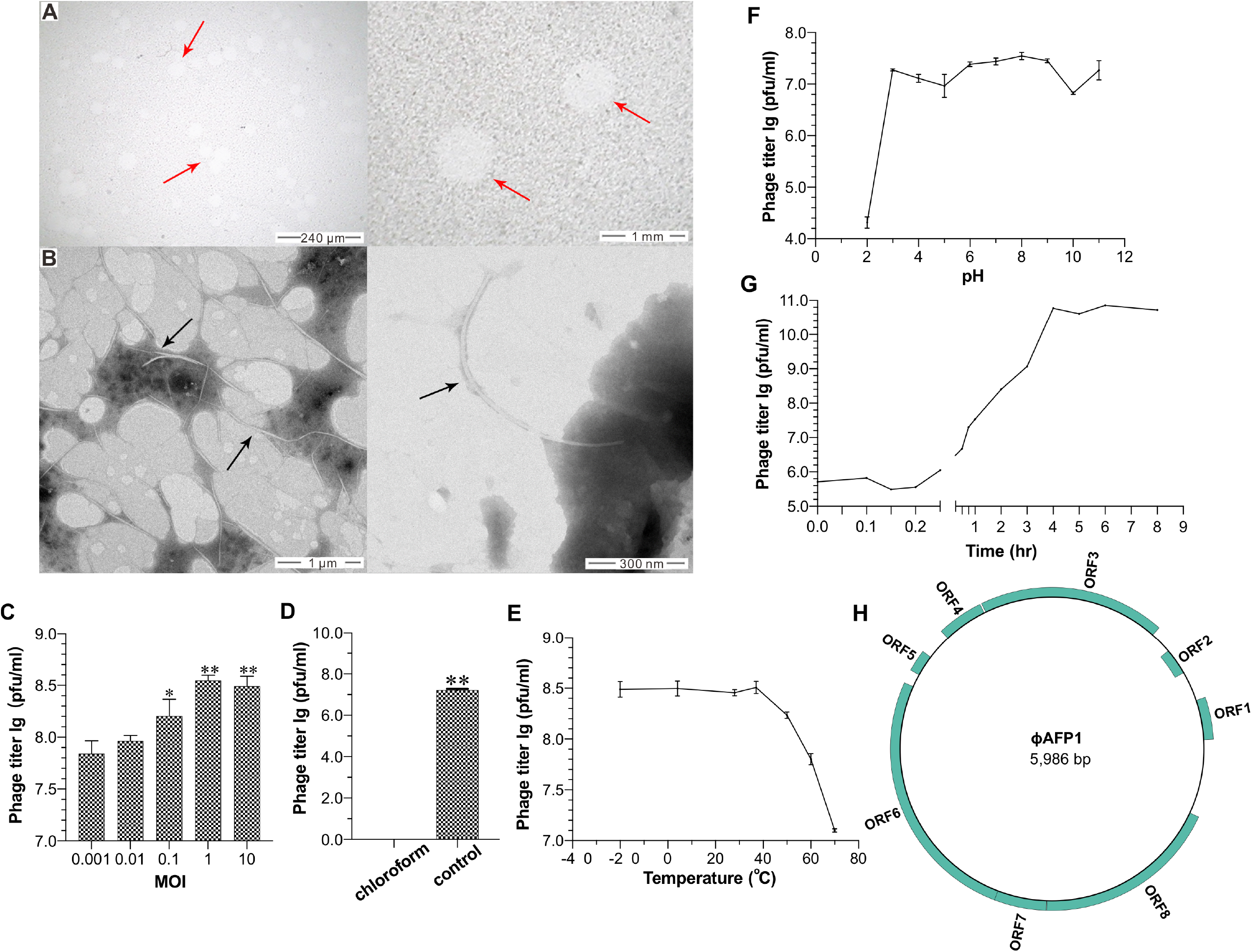
Plaque, morphology, and biological characterization of phage ϕAFP1. (A) The observations of plaques formed on the lawn of under different magnification. Red arrows indicated the plaques of phage ϕAFP1 against the lawn of host bacterium *A. abrolhosensis.* (B) Transmission electron microscopy observations of phage ϕAFP1 negatively stained. The black arrows indicated the phages. (C) Optimal multiplicity of infection (MOI) value of phage ϕAFP1. (D) Sensitivity of phage ϕAFP1 to Chloroform. (E) Biological stability of phage ϕAFP1 under different temperature conditions. (F) Sensitivity of phage ϕAFP1 to pH from 2 to 10. (G) One-step growth curve of phage ϕAFP1. The mean titer is shown from triplicate assays. The asterisk indicates significant differences between groups (* *p* < 0.05, ** *p* < 0.01) using One-way ANOVA. (H) Circular diagram of phage ϕAFP1 genome and predicted ORFs. Rects indicate ORFs in the plus or minus strand.

Multiplicity of infection (MOI) analysis was performed to determine the optimal MOI of ϕAFP1. The results showed that infection with phages at an MOI of 1 produced the highest phage titer (3.51×10^8^ pfu ml^-1^) (Figure 1C). The infectivity of phage ϕAFP1 was completely inactivated after treated with chloroform (Figure 1D). Phage ϕAFP1 was stable and retained nearly 100% infectious activity in a range from −20℃ to 37℃, but had a continuous and rapid decline in surviving phages titer when the temperature was higher than 37℃ (Figure 1E). The optimum pH range of ϕAFP1 was from pH 3 to 9, but had a substantial decline in phage titer with sharply decreased to 0.11% at pH 2 and 23.99% at pH 10, respectively (Figure 1F). The latent period of ϕAFP1 was approximately 12 min and then there was an ascent stage which continued for 228 min before entering the plateau period (Figure 1G). The burst size of ϕAFP1 presented that about 10 phage particles were produced per infected host cell. The agarose gel electrophoresis revealed that the genome size of ϕAFP1 is around 5-6 kbp (Supplementary Figure S1A). The genome was resistant to general restriction enzymes and RNase A but sensitive to DNase I and S1 nuclease (Supplementary Figure S1B and C), suggesting ϕAFP1 consists of ssDNA. The purified ϕAFP1 particles have a major protein band with theoretical molecular masses of ∼4 kDa visible on a Tris-tricine-SDS-PAGE (Supplementary Figure S1D).

To examine the host range of phage ϕAFP1, its infectivity was tested on five bacteria strains of different species/genera/families by spot test of diluted phage solution. The results showed that ϕAFP1 could infect none of the five bacteria strains (Supplementary Table 1). The association between ϕAFP1 and its bacterial host was further identified in sequence-based tools PHIS Detector, and there is not any predicted host found.

### Genomic analysis and annotation of ϕAFP1

The genome of phage ϕAFP1 composed of 5986 bp (GenBank accession no. MT975991with 40.34% G+C content, the completeness was estimated as 100% in CheckV. ϕAFP1 was predicted to encode eight open reading frames (ORFs) accounting for 82.69% of the genome, including three capsid proteins, two replication-related protein, a zonular occludens toxin (ZOT)-like protein and two proteins of unknown function (Figure 1H and Supplementary Table 2). ϕAFP1 contained a small amount of ORFs compared to other inoviruses (Supplementary Table 3).

Three-dimensional structures of ϕAFP1 ORFs were generated to predict the protein function using RoseTTAFold. Structural construction of the ORF4 model was obtained (Confidence: 0.82) (Figure 2A). Its structure was similar to the monomer of a homodimer single-stranded DNA binding protein (ssDBPs) (PDBID: 1PFS) encoded by the filamentous phages ϕPf3. The predicted protein consists of seven β-strands, the four-stranded antiparallel sheet (β1, β3, β4 and β5) across the molecule and two prominent hairpins are designated as the DNA binding wing and dyad loop respectively. The overall fold of structure was also partly similar to the monomer of ssDBP encoded by ϕPf3.

**Figure 2.**
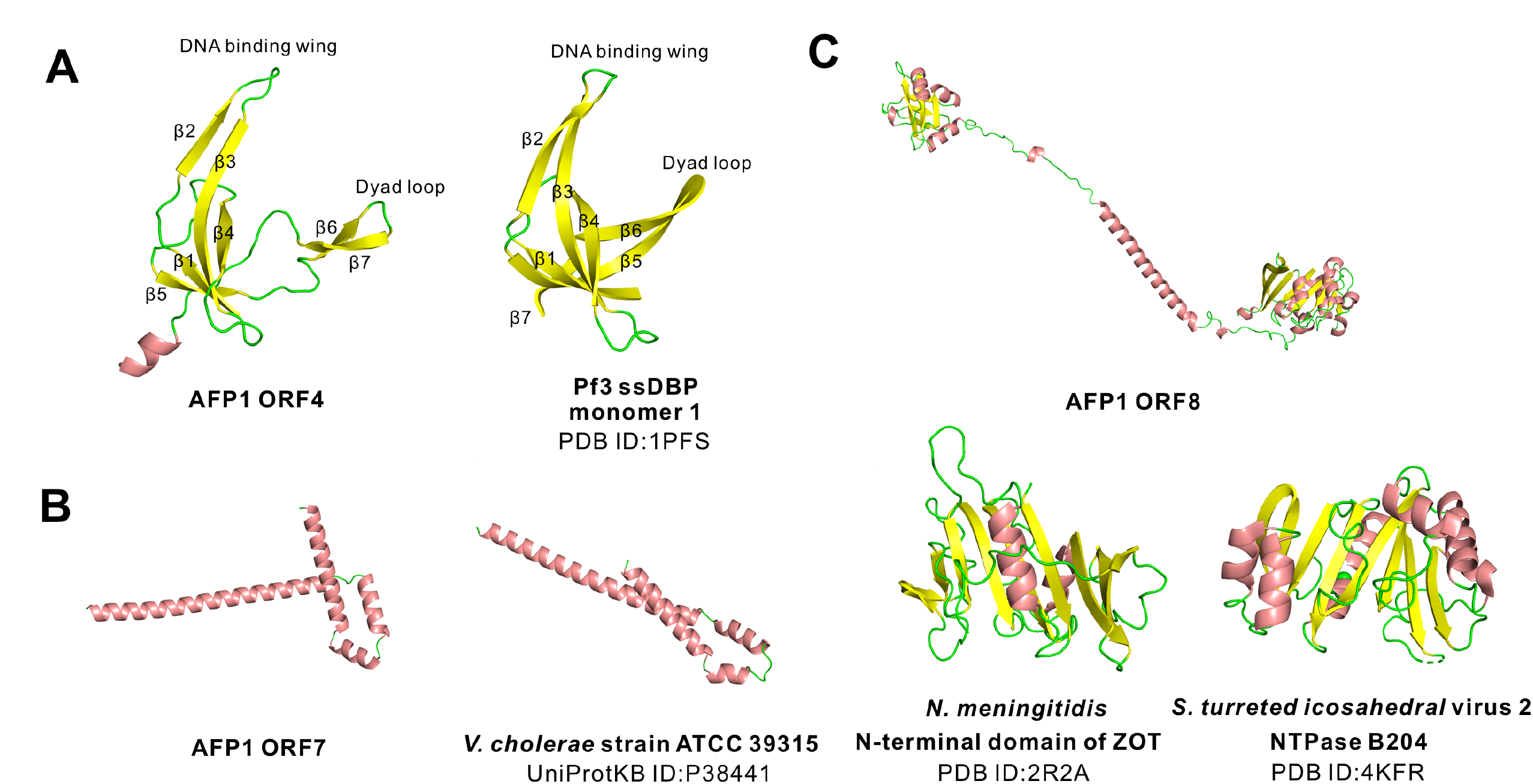
Predicted structures with highest confidence of ORFs proteins from ϕAFP1 and their homologous proteins. (A) ORF4 from ϕAFP1 and the ϕPf3 homologue (PDB: 1PFS, monomer 1). (B) ORF7 from ϕAFP1 and the *V. cholerae* strain ATCC 39315 homologue (UniProtKB ID: P38441). (C) ORF8 from ϕAFP1 and its homologues from the *N. meningitidis* N-terminal domain of ZOT (PDB: 2R2A) and *S. turreted icosahedral* virus 2 NTPase B204 (PDB: 4KFR).

The secondary structure of ORF3 (putative phage/plasmid replication protein) defines two domains, including OD (amino acids 1 to 265) and TD (amino acids 276 to 346) (Figure S2C). OD folds as an α-β sandwich with a central eight-antiparallel β-strands inserted with six helices flanking on two sides. TD is a three-helix bundle with one of the helices nearly perpendicular to the others.

The ORF7 contains a four-helix bundle with one of these helices nearly perpendicular to others (Confidence: 0.79) and is partly resemble to the protein of unknown function DUF2523 (Figure 2B), which is also identified as accessory cholera enterotoxin (P38441) in *Vibrio cholerae*, suggesting the putative role of ORF7 involving in the assembly of the ϕAFP1 virion at the membrane.

The ORF8 in ϕAFP1 encodes a predicted putative ZOT. The N-terminus of ORF8 adopts an αβα-fold, where eight-parallel β-strands (one is antiparallel) with six α-helices on one face and six α-helices on the opposite face, has a lesser extent similarities of the N-terminus of Zot from *Neisseria menigitidis* (Figure 2C). Furthermore, the considerable similarity of structural prediction of ORF8 to the P-loop containing nucleoside triphosphate hydrolase superfamily was observed (Figure 2C).

### ϕAFP1 represents a novel viral genus of *Inoviridae* and is a genetic mosaic

To investigate the evolutionary history of ϕAFP1, a phylogenetic tree was constructed based on the amino acid sequences of the whole complete genome from 36 *Inoviridae* and one *Paulinoviridae* (Supplementary Table 4). As shown in Figure 3A, ϕAFP1 was closely related to *Ralstonia* phage, but far from other characterized *Inoviridae* viruses. Network analysis of the ϕAFP1 genome together with the publicly available virus genomes showed that the virus sequence was disconnected from other viral clusters at the genus level (Supplementary Table 5). These results suggest that ϕAFP1 presents a new genus of *Inoviridae*.

**Figure 3.**
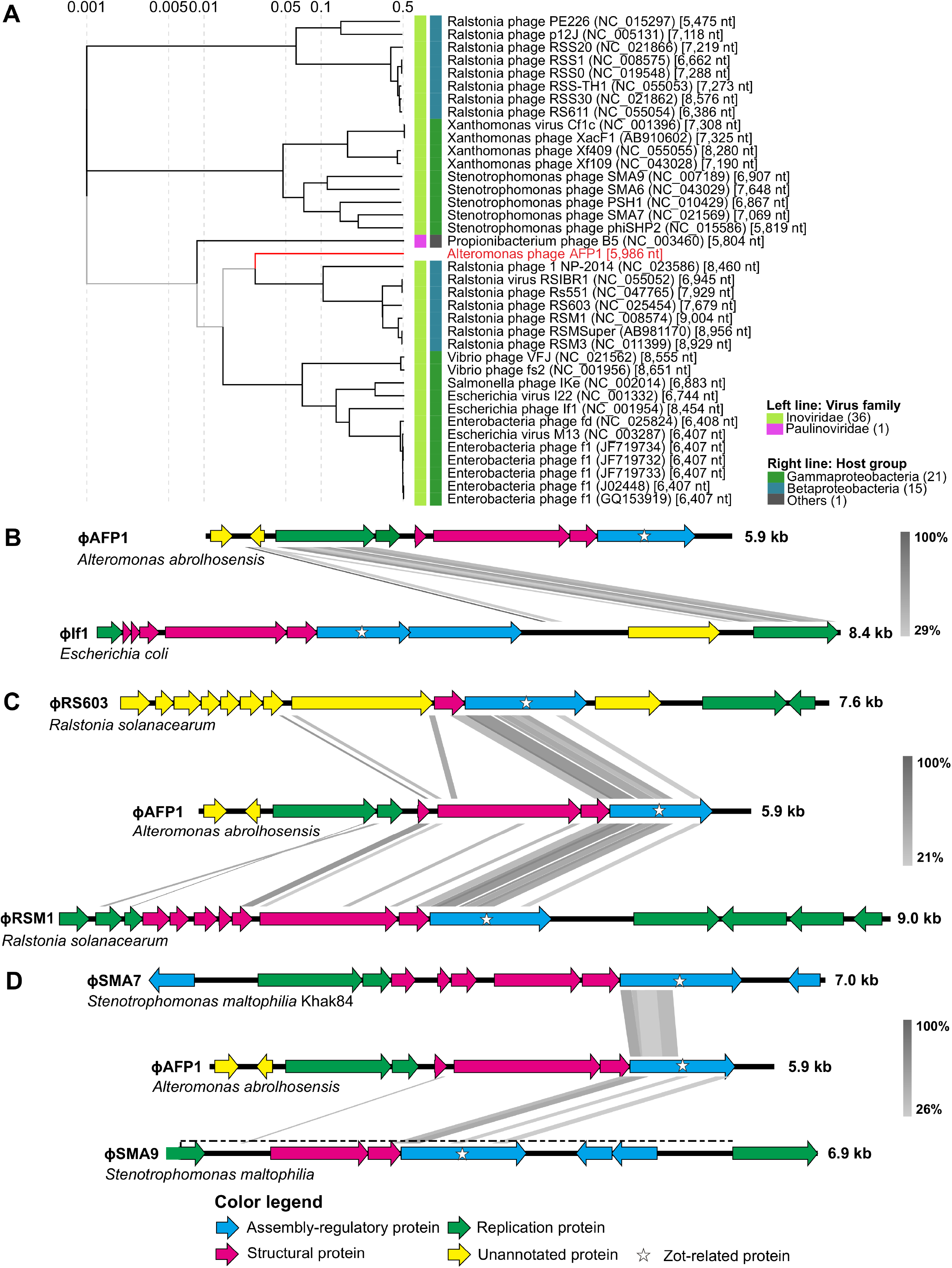
Phylogenetic trees and comparative analysis of ϕAFP1. (A) Phylogenetic tree generated based on amino acid sequences of complete genomic sequences of ϕAFP1 and other different filamentous phages in *Inoviridae* and *Paulinoviridae* family. Further details for phages represented on the tree were listed in **Supplementary Table 3**. (B, C, D) Comparative analysis of the linear genomic organization of ϕAFP1 and other three kinds of filamentous phages. Arrows with different colors represent the putative ORFs belonging to functional modules (Replication module, green; Structure module, red; Assembly-regulatory module, blue; unannotated genes, yellow). Further details for comparative analysis were listed in **Supplementary Table 2**.

To investigate the origin of ϕAFP1 and identify the function of its ORFs, a search for homologous sequences with other *Inoviridae* viruses was performed by tblastx (Supplementary Table 2). The genomic organization of phage ϕAFP1 was typically modular, including replication module, structural module, and assembly-regulatory module. ORF3 showed sequences similarity to the If1p10 of *Enterobacteria* phage If1 with the top hit (52.4% amino acid similarity) (Figure 3B), which was predicted to represent phage and plasmid replication proteins (gene II/X family). ORF4 had a certain level of similarity to ORF2 (ssDBP) of *Stenotrophomonas* phage, phiSMA6 (Supplementary Figure S3A). ORF5 had the highest sequences similarity with major capsid protein of *Ralstonia* phages (48.83%) (Figure 3C and Supplementary Figure S3B). ORF7 was aligned to the minor coat protein pVI of *Ralstonia* phages (Figure 3C and Supplementary Figure S3B). The amino acid sequence of ORF6 had 27.10%, 27.06% and 28.70% identity to the ORFs from *Ralstonia* phage RSM1, RSMSuper and RS603 (Figure 3C and Supplementary Figure S3B), respectively. Within the assembly-regulatory module, ORF8 of ϕAFP1 was predicted to encode the pI-like gene (the most conserved gene in inoviruses), the certain level of identity of ORF8 was found both in *Ralstonia* and *Stenotrophomonas* phages (Figure 3 and Supplementary Figure S3). The additional ORF1 and ORF2 had no significant similarity detectable to sequences in the databases. These results suggest that ϕAFP1 is genetic mosaics and the genes of mosaic structure most likely come from *Ralstonia* phage and *Stenotrophomonas* phage. Furthermore, the close relationships were verified by a phylogenetic tree based on the amino acid sequences of Zot-like proteins, showing that ϕAFP1 was more related to *Ralstonia* phage and *Stenotrophomonas* phage (Supplementary Figure S4).

### A temporally coordinated program of ϕAFP1 gene expression

To establish characteristic responses of the host infected by ϕAFP1, we firstly assembled the high-quality genome of *A. abrolhosensis* strain JKT1 (4 402 100 bp, GC 44.67%) from short and long sequencing reads with estimated completeness of 100% and contamination of 0.3%, and 3573/3699 genes were annotated in the bacterial genome (Supplementary Table 6).

Based on the one-step growth curve of ϕAFP1, approximately three-time points (6 min, 2 h, and 6 h) for latent, burst and plateau periods that spanned the entire infection cycle duration were selected as representative snapshots for exploring the gene expression patterns of both ϕAFP1 and host *A. abrolhosensis* by RNA-seq. The transcriptome of host *A. abrolhosensis* strain JKT1 during the phage infection cycle was progressively taken the place of phage ϕAFP1 transcripts, fluctuating from 0.0% at 6 min and peaking at 3.5% at 6 h (Supplementary Table 7), and the proportion of host reads reached a very high percentage around 96.3%∼99.7% (Figure 4A).

**Figure 4.**
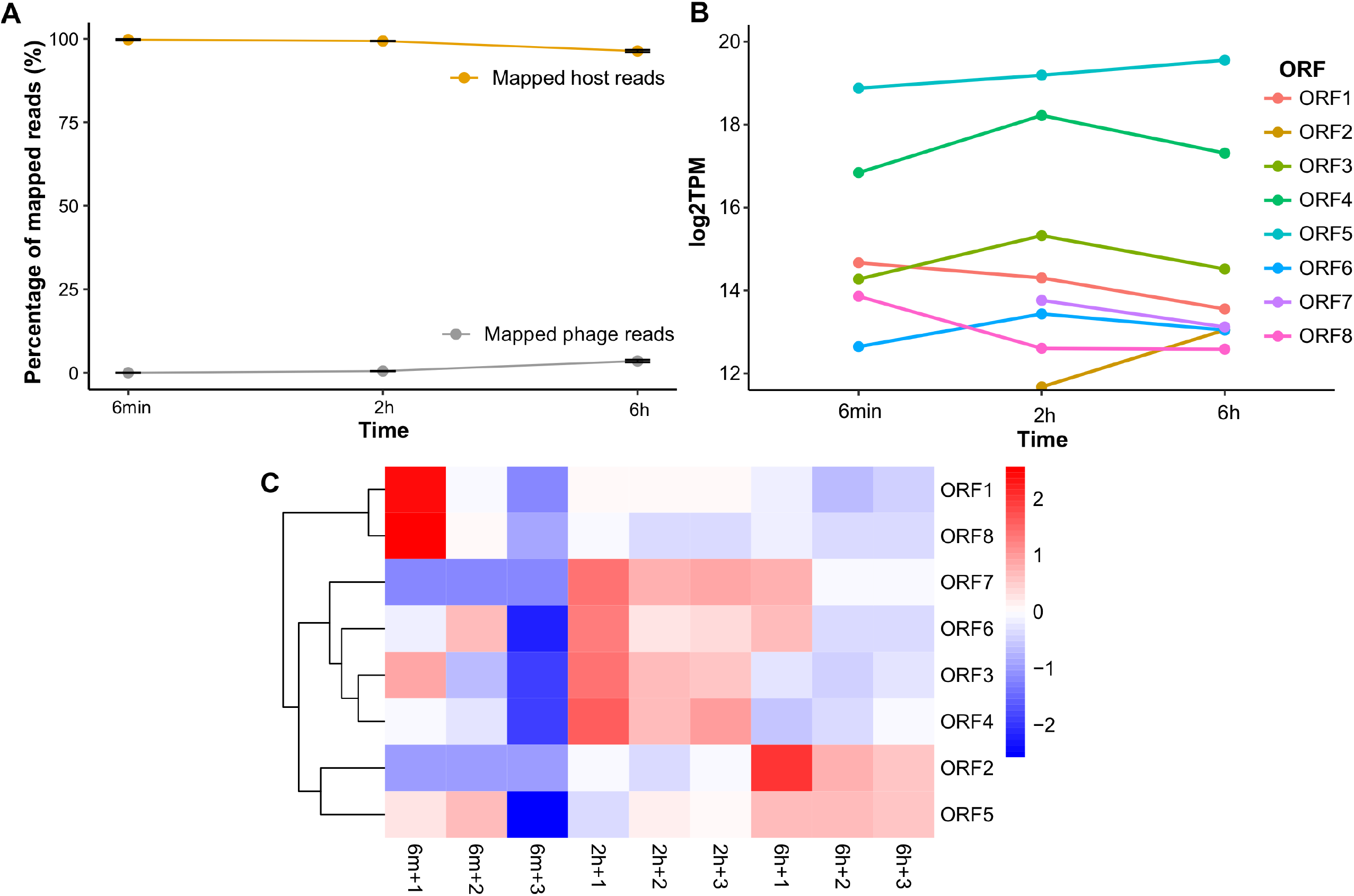
Temporal transcriptional profile of ϕAFP1. (A) Alignment of RNA read sets against the *A. abrolhosensis* (orange) or ϕAFP1 (grey) genome at three time points after infection. Data are displayed as the means SD from three independent experiments. (B) The relative abundance of the expression of phage ORFs at three time points. (C) Hierarchical cluster heat map of expression level of ϕAFP1 ORFs based on the TPM values of each gene. Further details for read alignment and expression abundance of phage ORFs can be found in **Supplementary Tables 7 and 8**.

According to the expression profiles of the individual genes (Figure 4B, Supplementary Table 8), ORF4 kept a relatively high expression level among all ϕAFP1 genes and represented the active replication of viruses in the development of phage infection. The temporally coordinated program of phage gene expression could be clustered into early, middle and late categories. ORF1 and ORF8 were highly expressed at 6 min and decrease thereafter, the abundance of ORF3, ORF4, ORF6 and ORF7 was at its peak at 2h, whereas genes expressed late were the ORF2 and ORF5 (Figure 4B and 4C).

### ϕAFP1 reprograms host metabolism and motility

To investigate the responses of host *A. abrolhosensis* to phage ϕAFP1 infection, we analyzed the differentially expressed genes (DEGs) and pathways in the host throughout the infection cycle. Totals of 0.87% (33/3779), 0% (0/3779), and 7.04% (266/3779) genes were differentially expressed at 6 min, 2 h, and 6 h in response to phage infection, respectively (Figure 5A and Supplementary Table 9). The differential fold change of DEGs measured by RT-qPCR was in good agreement with the RNA-seq data, including upregulated DEGs (*mmsB*, *prpB* and *phoB*) and downregulated DEGs (*fadA*, *flgE*, *flgL*, *flgG*, *hppD*, *prpC* and *galK*) (Supplementary Figure S5 and Supplementary Table 10).

**Figure 5.**
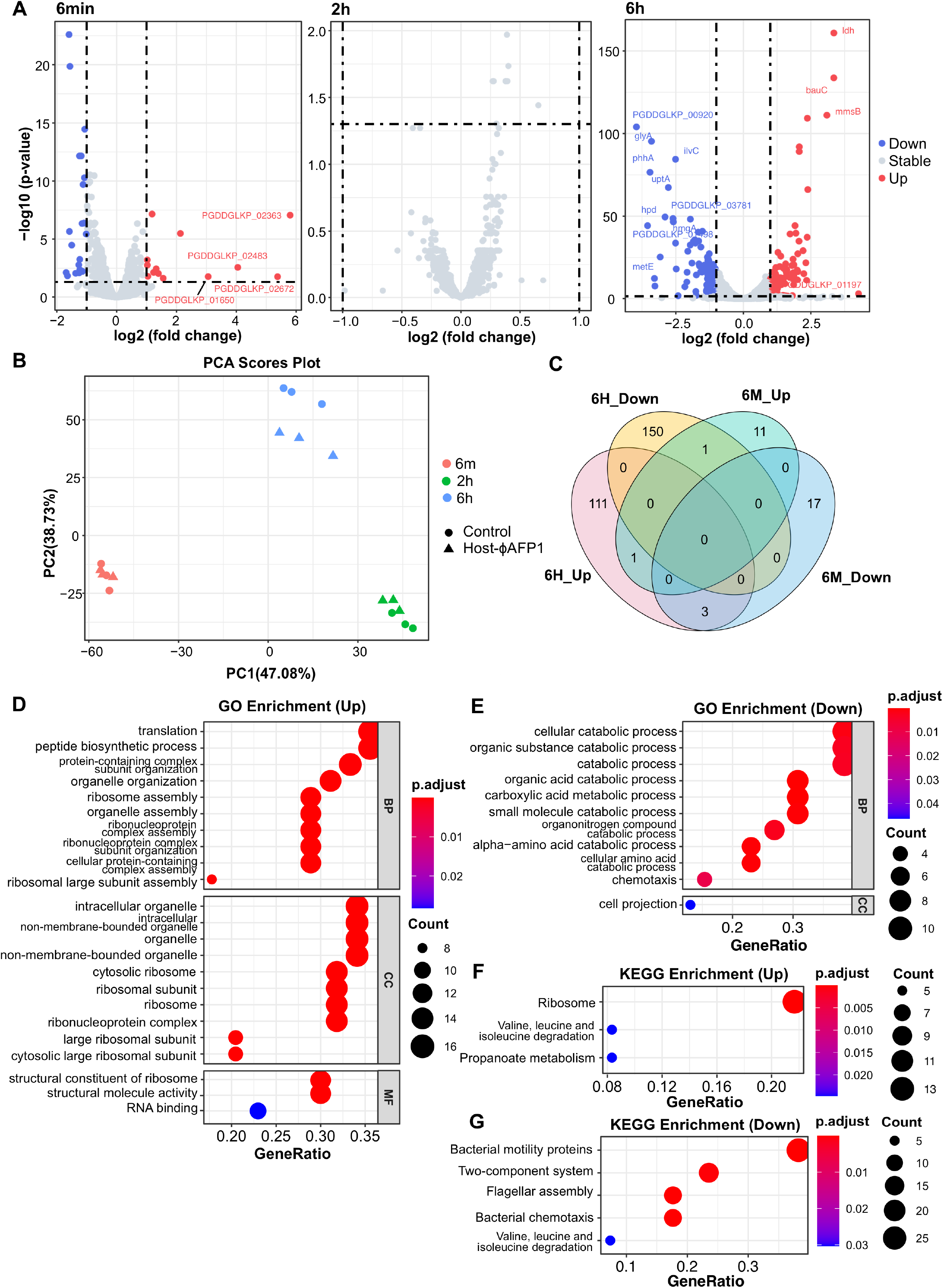
Impact of ϕAFP1 infection on its host transcriptome. (A) Volcano plot of the *A. abrolhosensis* transcriptome following phage infection compared with the uninfected control at three infection stages. (B) PCA plot of the expression of *A. abrolhosensis* genes in phage-infected or phage-uninfected groups. (C) Number and intersection of up- or downregulated DEGs at two infection stages. Significant enrichment GO analysis (D and E) and KEGG (F and G) categories of host DEGs (upregulated and downregulated genes) enriched at 6h after ϕAFP1 infection (*p*. adjust < 0.05). The enrichment *p*. adjust value of each pathway is shown as a color gradient. The number of genes enriched in each pathway is represented by the size of the points. Down, downregulated DEGs; Up, upregulated DEGs; Stable, non-significantly changed genes. Further details for DEGs of the host can be found in **Supplementary Tables 9**, **12 and 13**.

The expression of *A. abrolhosensis* genes changed greatly during ϕAFP1 infection, according to PCA (principal-component analysis) based on the TPM (transcript per million) values of host genes (Figure 5B). The most highly expressed genes at all the time points during infection encoded DNA binding and ribosomal-related proteins, which is varied among three groups (Supplementary Figure S6 and S7, Supplementary Table 11). Most of the DEGs were specific at 6 min and 6 h (except for upregulated DEG *cyoB*), and only four DEGs undulated between up- and downregulation (*hutD*, *prpB*, *yegQ* and gene encoding COG0517 FOG CBS domain) (Figure 5C). Moreover, the great majority of the DEGs were downregulated throughout the course of infection, indicating negative control by ϕAFP1 on suppressing host genes.

Ten DEGs (10/33) at 6 min were annotated in gene ontology (GO) and enriched in KEGG (Kyoto Encyclopedia of Genes and Genomes) pathways. Annotated upregulated genes were *gntX* and *cyoB*, while part of downregulated genes at 6 min encoded transport-related proteins (e.g. ExbD/TolR and BtuB), organic acid related metabolic proteins (e.g. PrpB, GalK, PrpC and HutU) (Supplementary Table 9). However, our result showed that none of the differentially expressed genes were found at 2 h post ϕAFP1 phage infection (the burst period) compared to the uninfected host (Supplementary Table 9).

To gain the biological functions of the significantly changing host genes, 83/266 DEGs at 6 h were annotated with GO function by Eggnog-mapper, showing the remarkable function classifications of annotated DEGs (Figure 5D and E, Supplementary Table 12). Most upregulated genes of the 6 h group are enriched in translation, ribosomal structure, ribosome and organelle biogenesis and transcription, revealing that ϕAFP1 highly demanded the translation machinery of bacterial host. The majority of the downregulated genes are involved in the pathways of chemotaxis and organic substance catabolic process (3.4-fold change gene *phhA*, 2.6-fold change gene *hmgA*), including organic acid, small molecule, organonitrogen and amino acid.

62.4% (166/266) of the DEGs at 6 h were clustered as 68 KEGG pathways, the pathway categories of up- and downregulated DEGs were marked respectively (Figure 5F and G, Supplementary Table 13). The significantly changed upregulation pathways were ribosome, valine, leucine and isoleucine degradation and propanoate metabolism, but the pathways bacterial motility proteins, two-component system, flagellar assembly, bacterial chemotaxis, valine, leucine and isoleucine degradation were downregulated. The DEGs involved in valine, leucine and isoleucine degradation pathway participated in different metabolic processes, such as the downregulated formation (*ilvE*) and upregulated degradation (*pdhA*) of methyl-2-oxobutanoate (Supplementary Figure S8). A total of 27 downregulated DEGs in *A. abrolhosensis* strain JKT1 were related to bacterial motility pathways, including genes related to flagellar assembly (e.g. *fla, flg*, *fli* family) (Supplementary Figure S9), pilus assembly protein (PilY1 and PilX), and chemotaxis genes (*cheA, cheW*, *cheY*, and *aer2*) (Supplementary Figure S10), suggesting the reduction of flagella dependent cell movement caused by ϕAFP1. Functional analysis mentioned above reveals the step-by-step control of ϕAFP1 on host genes.

## Discussion

In this study, we report the isolation and complete genome sequence of a filamentous phage designated ϕAFP1 from the South China Sea using the *A. abrolhosensis* strain JKT1 as the host bacterium. Until now, only ten published reports about bacteriophages isolated from *Alteromonas* are available, including *Corticoviridae* phage PM2 (50), five *Siphoviridae* phages PB15, JH01, P24, XX1924, and R7M (30, 51–53), *Myoviridae* phage V22 (54), *Podoviridiae* phages altaD45 P1 to P4 (55), *Autographiviridae* phages H4/4’ (56) and R8W (57), and *Mareflavirus* phage ZP6 (58). Different from the above phages, ϕAFP1 consisted of flexible filament (width=10 nm, length=900 nm), suggesting that this phage was a member of the family *Inoviridae*. To date, only two filamentous phages infecting bacteria of the family *Alteromonadaceae* have been isolated and reported, including phage f327 infecting *Pseudoalteromonas* (59) and SW1 infecting *Shewanella* (60). To the best of our knowledge, this is the first report of the filamentous phage infecting *Alteromonas*. The family *Inoviridae* includes 21 genera and 27 species with genomes of about 5.5-10.6 kb encoding 7-15 proteins (8). ϕAFP1 contains fewer coding genes (8 ORFs) and has the small genome size (5.9 kb). The genus demarcation of inoviruses is set at 50% amino acid identity on the pI-like gene (i.e. Zot) and the major capsid protein, and there is no significant similarity of both Zot and major capsid proteins found between ϕAFP1 and other inoviruses. In addition, the phylogenetic analysis and gene-sharing network suggested that ϕAFP1 represents a new viral genus of *Inoviridae*.

One of the most evident features of bacteriophage genomes is their genetic mosaicism (61). Previous studies have observed mosaicism within several phages, such as *Pseudomonas* phages (62–64), *Actinobacteria* phages (65) and *Mycobacteria* phages (66). Phage mosaic genomes are relatively common in temperate phages, revealing the horizontal exchange of genetic material between distinct phages and phages or their hosts. Inoviruses genomes are often shaped by frequent recombination and a mosaic of genes emerged as a result (67–69), which plays a key role in the evolution of these viruses by leading to the acquisition of non-orthologous genes or gene exchange among highly divergent orthologs (70, 71). Here, comparative genomic analysis demonstrated the mosaic nature of ϕAFP1 genome, which is closely related to *Ralstonia* phages and *Stenotrophomonas* phages. The genes encoding Zot-like protein in ϕAFP1 are similar to both these phages, while the structure-related genes likely were contributed by *Ralstonia* phages. Genetic exchange may yield new combinations of genes and protein domains or give rise to non-functional genomic trash (72). These exchange events among ϕAFP1, *Ralstonia* phages and *Stenotrophomonas* phages result in the inclusion of key genes of inovirus genomic modules, we speculate that there might be frequent communication between *Ralstonia* phages and *Stenotrophomonas* phages. Although we did not identify novel hosts of ϕAFP1 in this study, considering that genetic recombination more likely occurs between phages whose host ranges are more similar, we predicted that the hosts of *Ralstonia* phages and *Stenotrophomonas* phages might also be the potential hosts of ϕAFP1.

The transcriptional results manifest that the takeover and shutoff of *A. abrolhosensis* gene expression by ϕAFP1 was step-by-step but not suddenly within the long-term associations with bacterial. In general, early expressed genes of ϕAFP1 peaked in the early stages of infection and altered the expression of hosts genes associated with putative bacterial defense responses and phage hijacking mechanisms. Similarly, this phenomenon has also been reported in ϕAbp1 infected AB1 (19). Next, during the middle stage of infection (2 h), all ϕAFP1 middle genes expressed but number of host DEGs returns to zero. This may be due to the chronic infection of ϕAFP1. Finally, a balance of phage interactions with host was achieved at the plateau stage (6 h).

The lysogenic phage pattern of ϕAFP1 exerted the step-by-step control on host genes, especially on genes belonging to DNA, RNA, and protein biosynthesis, central metabolism and bacterial motility. The upregulation of translation processes related DEGs at 6 h indicated ϕAFP1 hijacked cellular resources for the assembly of phages. The findings that phages hijacked synthesis systems of the hosts were also observed in other phages (19, 73, 74). In contrast, it was reported that phage infection could stop the host macromolecular synthesis promptly and completely (75). Pioneering work on bacterial motility revealed that phages could exert multiple roles in the hosts, including contributing to searching more adaptive environments, serving as virulence factors for pathogens and being conducive to symbiotic relationships (76–79). The downregulation of host genes related to material catabolic processes and bacterial motility-related pathways was consistent with other temperate phage infections (18, 23, 80, 81). While filamentous phage f327 was reported to improve the motility of the host in Arctic Sea ice (22). Considering that there is a large amount of energy cost for flagellar biosynthesis, bacterial motility, and chemotaxis for the host, thus downregulating the key genes of these processes may be useful for ϕAFP1 utilizing cellular resources.

There is a range of responses in different phage-bacterium systems, and the trend of expression changes depends on the host-phage interactions (82). Our global analysis of host regulation indicated that the infection of ϕAFP1 did not interfere with host regulation at the burst period, which may be the exception rather than the rule known so well for phage infection. It was observed that ϕAFP1 exerted a temperate infection on *A. abrolhosensis*, performed with a long release period, relatively low burst size (10 pfu cell^-1^), quite minor proportions of phage transcripts and some upregulated host material metabolism processes to promote the use of biomolecules. The small and simple genome of ϕAFP1, as well as its unique chronic infection may cause the distinctive shifts in expression of host genes during the infection, however, the precise mechanism of ϕAFP1 infection needs to be further studied.

To assess the global diversity of ϕAFP1, the ecological distribution of ϕAFP1 was characterized. However, there was no hit found in all selected dataset, this is mainly because the database representation for sequence comparisons of ssDNA viruses is so poor. The ssDNA viruses, albeit extremely abundant, viral metagenomes of them were not fully collected, while dsDNA phages account for 90% of the phages reported in the literature (70, 83).

In conclusion, novel *Inoviridae* phage ϕAFP1 isolated from the South China Sea was the first report of the filamentous phage infecting *Alteromonas*. The biological and genomic characterization of ϕAFP1 could provide new information for marine virology. Furthermore, we found diverse and novel expression patterns of phage and host at different time points during infection, which reveal reprogramming of the host metabolism and repression of the bacterial motility system. Our results revealed a formerly unidentified regulatory tactics that diverges from the phage infection paradigm and contributes to a broader understanding of phage-bacterium interactions.

## Acknowledgements

The work was supported by the National Natural Science Foundation of China (32173001). We thank Dr. Simon Roux and Dr Zengpeng Li for the helpful discussions on our analyses.

---

**Table S1 Analysis of the host range of phage ϕAFP1.**

**Table S2 Comparative genomic analysis of ϕAFP1 and other filamentous phages based on amino acid identity.**

**Table S3 The summary of some inovirus isolates.**

**Table S4 Accession numbers of sequences used in phylogenetic trees construction.**

**Table S5 Overview of viral clusters by vConTACT2 for ϕAFP1.**

**Table S6 EggNOG annotation results for Alteromonas abrolhosensis.**

**Table S7 Alignment of RNA read sets against the A. abrolhosensis and ϕAFP1 genome at three time points after infection.**

**Table S8 Expression abundance of phage ORFs at three time points after infection.**

**Table S9 Differentially expressed Analysis of host genes at three time points after infection.**

**Table S10 Relative expression of host genes measured by RT-qPCR and RNA-Seq.**

**Table S11 Expression abundance of host genes at three time points after infection.**

**Table S12 GO analysis of host DEGs (up- and downregulated genes) enriched at 6h after ϕAFP1 infection.**

**Table S13 KEGG categories of host DEGs (up- and downregulated genes) enriched at 6h after ϕAFP1 infection.**

**Table S14 Primers used for qRT-PCR.**

**Figure S1 Genomic DNA and protein analysis of phage ϕAFP1.** (A) Agarose gel electrophoresis of the genomic DNA of phage ϕAFP1. Lane 1, phage ϕAFP1 genomic DNA. (B) Agarose gel electrophoresis of the genomic DNA of phage ϕAFP1 treated with DNase I (Lane 1), S1 nuclease (Lane 2) or RNase A (Lane 3). Lane 4: untreated ϕAFP1 genomic DNA. (C) Agarose gel electrophoresis of the genomic DNA of ϕAFP1 treated with general restriction enzymes, EcoR Ⅰ(Lane 1), Pst Ⅰ(Lane 2), Xba Ⅰ(Lane 3), BamH Ⅰ(Lane 4), Hind Ⅲ (Lane 5) or untreated (Lane 6). (D) Tris-tricine-SDS-PAGE gel image of purified phage ϕAFP1 staining with Coomassie brilliant blue R-250. Lane M, 1 kbp DNA Marker.

**Figure S2 Predicted structures with highest confidence of ORFs proteins from ϕAFP1.** ORF1 (A), ORF2 (B), ORF3 (C), ORF5 (D) and ORF6 (E) from ϕAFP1.

**Figure S3 Comparative analysis of the linear genomic organization of ϕAFP1 and other filamentous phages, Stenotrophomonas phage (A) and Ralstonia phage (B).** Arrows with different colors represent the putative ORFs belonging to functional modules. Genes belonging to the replication module are marked in green, genes belonging to the structure module are marked in red, genes belonging to the assembly-regulatory module are marked in blue, and unannotated genes are marked in yellow. Further details for comparative analysis were listed in **Supplementary Table 2**.

**Figure S4 Maximum-likelihood phylogenetic trees generated based on amino acid sequences of Zot-like proteins (pI) of ϕAFP1 and other different filamentous phages.** NCBI GenBank accession numbers of phages represented on the tree were listed. The numbers at the nodes represent Bootstrap values and scale bars indicate the average number of substitutions per site. Further details for phages represented on the tree were listed in **Supplementary Table 3**.

**Figure S5 RNA-seq data was verified by RT-qPCR.** Comparison of selected DEGs from RNA-Seq and RT-qPCR based on relative expression (A) and Linear regressions (B). Further details for RT-qPCR can be found in **Supplementary Table 10 and 14**.

**Figure S6 Distance heat map among different samples based on ϕAFP1 genes (A) and *A. abrolhosensis* genes (B) using bray-curtis methods.**

**Figure S7 Clustering heat map of the top 30 expressed *A. abrolhosensis* genes among different samples.** Further details for expression abundance of host genes can be found in **Supplementary Table 11**.

**Figure S8 KEGG pathway diagram of valine, leucine and isoleucine degradation (map00280).** The red border indicates up-regulation of differentially expressed genes and blue border indicates down-regulation.

**Figure S9 KEGG pathway diagram of flagella assembly (map02040).** The blue border indicates down-regulation of differentially expressed genes.

**Figure S10 KEGG pathway diagram of bacterial chemotaxis (map02030).** The blue border indicates down-regulation of differentially expressed genes and some genes of A. abrolhosensis clustered into map02030 are shown in the diagram.

